# AutoRELACS: Automated Generation And Analysis Of Ultra-parallel ChIP-seq

**DOI:** 10.1101/2020.03.30.016287

**Authors:** L. Arrigoni, F. Ferrari, J. Weller, C. Bella, U. Bönisch, T. Manke

**Affiliations:** Max Planck Institute of Immunobiology and Epigenetics, Freiburg, Germany; Faculty of Biology, University of Freiburg, Freiburg, Germany

## Abstract

Chromatin immunoprecipitation followed by sequencing (ChIP-seq) is a method used to profile protein-DNA interactions genome-wide. RELACS (Restriction Enzyme-based Labeling of Chromatin *in Situ*) is a recently developed ChIP-seq protocol that deploys a chromatin barcoding strategy to enable standardized and high-throughput generation of ChIP-seq data. The manual implementation of RELACS is constrained by human processivity in both data generation and data analysis. To overcome these limitations, we have developed AutoRELACS, an automated implementation of the RELACS protocol using the liquid handler Biomek i7 workstation. We match the unprecedented processivity in data generation allowed by AutoRELACS with the automated computation pipelines offered by snakePipes. In doing so, we build a continuous workflow that streamlines epigenetic profiling, from sample collection to biological interpretation. Here, we show that AutoRELACS successfully automates chromatin barcode integration, and is able to generate high-quality ChIP-seq data comparable with the standards of the manual protocol, also for limited amounts of biological samples.

## BACKGROUND

Chromatin immunoprecipitation followed by sequencing (ChIP-seq) is a widely used method to study protein-DNA interactions genome-wide (1). Despite the enormous contribution that ChIP-seq has brought to our understanding of epigenetic and transcriptional control, the traditional ChIP-seq protocol (2,3) presents various limitations. For example, it requires substantial amounts of biological input material, which is often a limiting factor in relevant clinical settings, and it is low-throughput, which prevents comprehensive epigenetic profiling. Furthermore, the protocol is poorly standardized across cell types, resulting in a high degree of technical variability that hampers biological interpretation of the data.

Over the last ten years, much work has been devoted to address these and other shortcomings (4–8). In line with these efforts, we have recently developed RELACS (Restriction Enzyme-based Labeling of Chromatin *in Situ*), a method that employs chromatin barcoding to enable high-throughput generation of ChIP-seq experiments (9). RELACS works reliably with low input material and can be used for quantitative ChIP-seq analysis (9,10). The method is highly standardized, and could potentially be scaled to profile hundreds of samples in parallel for tens of DNA-binding proteins at once. Yet, the current manual implementation is limited by human processivity in both data generation and data analysis.

To match the ideal potential of this methodology, we have implemented an automated version of the RELACS protocol, named AutoRELACS, using the liquid handler Biomek i7 automated workstation (Beckman Coulter). While other automated ChIP-seq implementations already exist (11,12), they still require a large amount of sample material, and they do not utilize the enormous multiplexing potential of barcoded chromatin. The scope of these methods is limited to data generation and lack an integrated bioinformatics workflow that streamlines standard computational tasks (e.g. QC, DNA-mapping, peak calling). AutoRELACS, on the other hand, couples the high-throughput generation of ChIP-seq data with the scalable and modular computational pipelines offered by snakePipes (13). From version 1.2.3, snakePipes’ DNA-mapping routine can handle RELACS data by performing demultiplexing of fastq files on RELACS adaptors and UMI-based deduplication. Together, AutoRELACS and snakePipes build a continuous workflow that automates ChIP-seq data generation and analysis, allowing for unprecedented processivity.

In this work, we test the performance of AutoRELACS by assessing 1) the scalability of the chromatin barcode integration step, 2) the quality of the generated data in comparison to the benchmark set by the manual protocol, and 3) the sensitivity of the automated method when working with low (≤ 25.000 cells/sample) and very low (≤ 5.000 cells/sample) cell numbers. We show that AutoRELACS is a scalable method that can generate high quality ChIP-seq data, comparable with the standards of the manual protocol. We finally show that AutoRELACS provides reliable epigenetic profiling also with limited input biological material.

## MAIN

### AutoRELACS is a scalable method for the generation and analysis of ultra-parallelized ChIP-seq data

RELACS (Restriction Enzyme-based Labeling of Chromatin *in Situ*) is a method that enables the high-throughput generation of ChIP-seq experiments (9). To increase the standardization and the scalability of this approach, we have developed AutoRELACS, an automated implementatin of the RELACS protocol using the liquid handler Biomek i7.

The AutoRELACS workflow is conceptually divided in six parts: four fully automated (A) processes intermitted by two manual (M) steps (Fig 1a). First, cells are manually processed to isolate the nuclei (14) and to digest the chromatin within the nuclear envelope (step 1 - M). Next, using the liquid handler Biomek i7, the chromatin from each sample is barcoded and pooled into a unique masterbatch (step 2 - A). Using focused sonication, nuclei are lysed and the barcoded chromatin is released (step 3 - M). The final three steps of the protocol have been fully automated and require minimal human supervision. These include the chromatin immunoprecipitation (ChIP) reactions and washing steps of beads-bound immunocomplexes (step 4 - A), decrosslinking, DNA purification and PCR amplification (step 5 - A) and, after sequencing, barcode demultiplexing and bioinformatics analysis with snakePipes (step 6 -A) (13).

**Figure 1:**
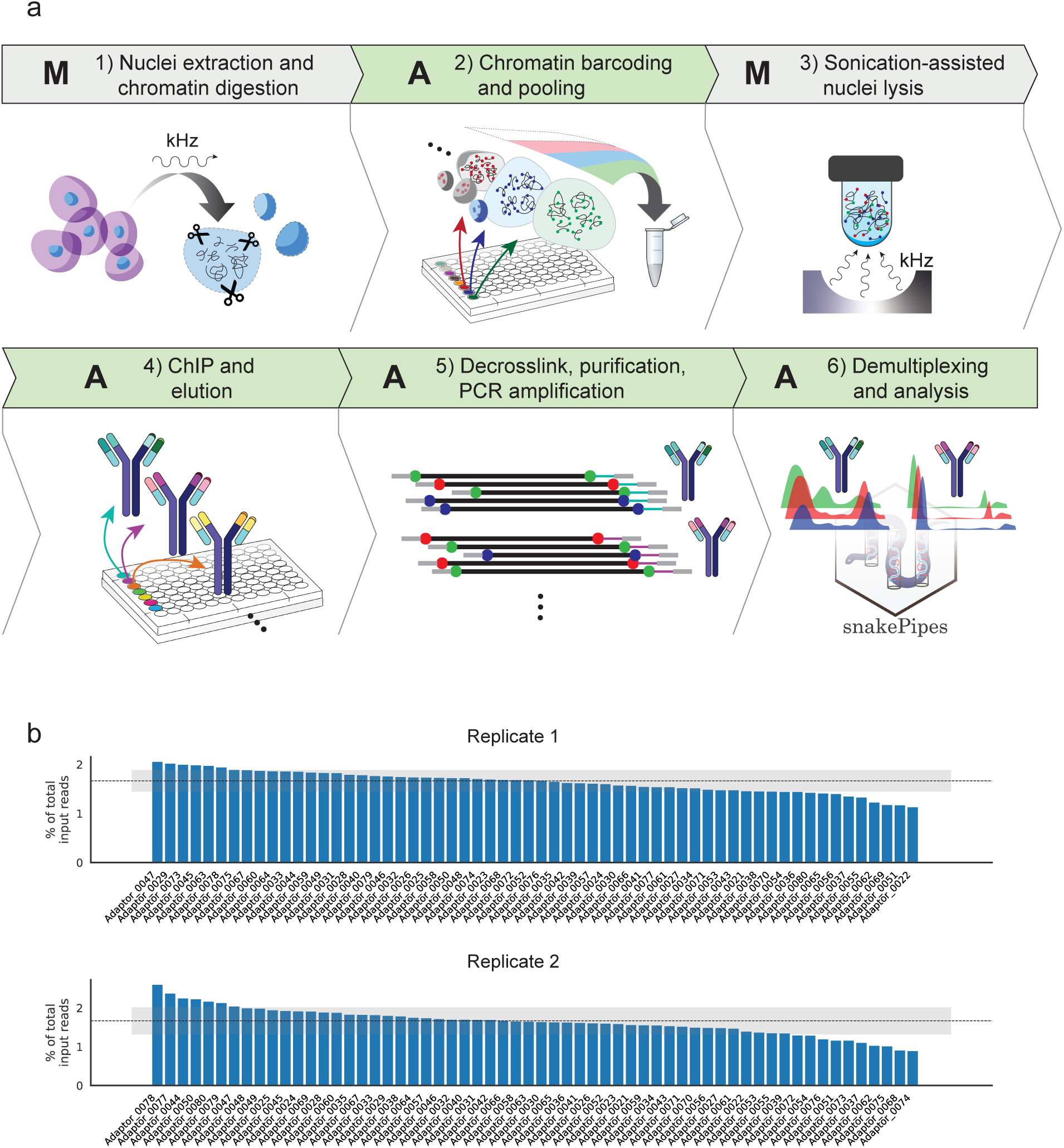
AutoRELACS workflow ensures comprehensive integration of RELACS barcodes. a) Overview of AutoRELACS protocol. 1-M) Nuclei of formaldehyde-fixed cells are extracted manually using adjusted ultrasound (14). The nuclear envelope is permeabilized, and the chromatin digested *in situ* using a 4-cutter restriction enzyme (RE). 2-A) Digested chromatin from each sample is automatically barcoded. Upon completion, the liquid handler pools all barcoded samples into a unique tube (Biomek i7 program: “RELACS_Barcoding”). 3-M) Pooled samples are collected by the user and nuclei are lysed using focused sonication. 4-A) The barcoded chromatin is aliquoted according to the number of required immunoprecipitation (IP) reactions into corresponding ChIP reaction mixes. The ChIP reactions are carried out overnight in parallel at room temperature on the Biomek i7 workstation. Upon completion, the ChIP-ped chromatin is sequestrated using beads and automatically washed 4 times at increasing stringency conditions and finally eluted in the elution buffer (Biomek program: “RELACS_ChIP_Elution”). 5-A) Subsequently, the eluted chromatin is decrosslinked and the DNA is purified. DNA is amplified via PCR using primers carrying Illumina dual indexes. Optionally, the liquid handler performs multiple rounds of purification and size selection using Ampure XP beads (Biomek program: “RELACS_Decrosslink_FinalLibraries”). A: Automated; M: Manual. 6-A) Libraries are sequenced on Illumina’s sequencing devices. Upon completion of the sequencing run, bcl2 files are automatically converted to fastq format and input into the fully automated ChIP-seq workflow available as part of the snakePipes suite (13). SnakePipes’ ChIP-seq workflow performs demultiplexing of reads on RELACS custom barcodes, quality controls, mapping and filtering of duplicate reads using unique molecular identifiers (UMI), and further downstream analysis like generation of input-normalized coverage tracks and peak calling. b) Distribution of RELACS barcodes in two independent input chromatin pools. 60 barcodes are integrated into the digested chromatin of two independent batches of S2 cells. Sequencing of the input chromatin pool for replicate 1 (upper panel) and replicate 2 (lower panel), reveals the percentage of input reads for each barcode used (y-axis). The ideal uniform distribution (100/60) is represented as a dotted line. The shaded gray area shows one standard deviation from the mean of the observed distribution.

The integration of sample-specific RELACS barcodes into the digested chromatin (Fig 1a, step 2) is key to the success of the method. To test the performance of automated and parallelized RELACS barcode integration, 60 custom barcodes were designed, each composed of a 4 nucleotide (nt)-long unique molecular identifier (UMI), followed by a 8 nt-long barcode with 50% GC content (note that after combining forward and reverse reads, each fragment is tagged by a 8-nt long UMI). These adaptors were used to label the chromatin of 60 batches of S2 cells (*Drosophila melanogaster*) in duplicates using the Biomek i7 workstation (Fig 1b). Results show that all barcodes are present within the pooled chromatin in both replicates, with a distribution of barcode representation equal to 1.64% ± 0.22% and 1.64% ± 0.35% for replicate 1 and 2 respectively, close to the uniform expectation of 1.667 % (Fig 1b, dashed line).

In summary, we show that AutoRELACS can be used to uniformly integrate multiple barcodes in a fully automated fashion, allowing for ultra-parallelized processing of a considerable number of samples in one single run.

### The quality of AutoRELACS ChIP-seq data is comparable with manual RELACS

Next, we test the quality of the ChIP-seq data generated with AutoRELACS and we compare it with the results from the previously published manual RELACS protocol. To this end, we run in parallel a manual and an automated RELACS experiment where we digest and barcode 28 batches of S2 cells and we immunoprecipitate against H3K4me3, H3K27ac and H3K27me3.

The histone modification profiles generated with manual RELACS and with AutoRELACS are overall similar. The variance present in the first two principal components of the normalized coverage matrix (computed on the merged peaks set) discriminates between the three histone modifications, regardless of the method used (Fig 2a). Comparison of the metaprofiles of the merged scores over peaks shows identical signal for H3K4me3, while H3K27ac and H3K27me3 present a slightly lower median coverage in AutoRELACS compared to the manual procedure (Fig 2b). Nevertheless, these differences do not impinge on the sensitivity of the assay. Visual inspection of the normalized coverage reveals high similarity between the two RELACS implementations (Fig 2c), while high overlap (80-90%) is observed between the peaks called in the two datasets (Fig S1a).

**Figure 2:**
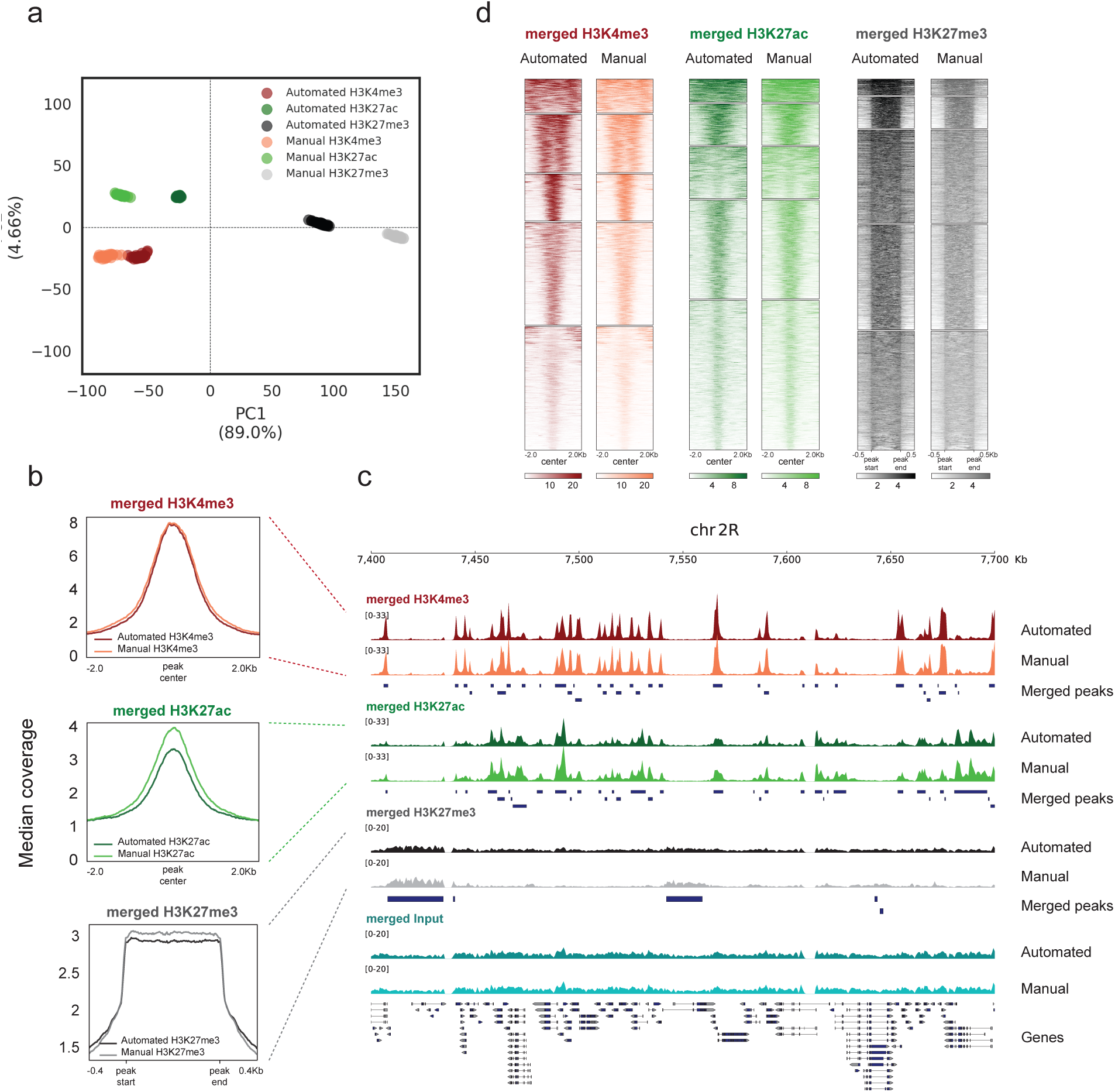
AutoRELACS ChIP-seq data are comparable with the standards of the manual protocol. a) Principal component analysis (PCA) of the normalized coverage matrix computed on the merged peak set between H3K4me3, H3K27ac and H3K27me3, as generated by AutoRELACS (Automated) and manual RELACS (Manual). For each mark and protocol implementation, all 28 demultiplexed technical replicates are shown. The 10000 most variable loci across all marks are input into the PCA. b) Metaprofile of the median normalized coverage computed over H3K4me3 (upper panel), H3K27ac (central panel) and H3K27me3 (lower panel) peaks. Each panel shows the signal generated with AutoRELACS and manual RELACS from a merge of all 28 technical replicates. c) Data tracks of the merged signal of the 28 technical replicates for H3K4me3 (red), H3K27ac (green), H3K27me3 (grey) and Input (cyan) on the dm6 locus chr2R:7,400,000-7,700,000. For each mark, we show the profile generated by AutoRELACS (Automated) and manual RELACS (Manual) and the merged set of peaks called in the two datasets (Merged Peaks). d) Heatmaps showing the clustered signal (k=5) on a merged set of peaks, as identified in the AutoRELACS (Automated) and in the manual RELACS (Manual) dataset, for H3K4me3 (left panel), H3K27ac (central panel) and H3K27me3 (right panel). The similarity between each pair of tracks indicates that there are no obvious implementation-specific biases.

To provide a global overview for all enriched regions, we cluster (k=5) the signal of H3K4me3, H3K27ac and H3K27me3 using the manual and the automated RELACS data on a common merged peaks set (Fig 2d). We do not observe any set of peaks that are specific to manual RELACS or AutoRELACS, which shows no obvious implementation-specific biases.

Together, we show that AutoRELACS yields high quality ChIP-seq data that are overall comparable with the manual RELACS protocol.

### AutoRELACS works reliably with low cell numbers

RELACS can generate robust epigenetic profiling with low cell numbers (9). To test the sensitivity limits of AutoRELACS, we barcode 4 batches of HepG2 cells and we aliquote the chromatin into two pools containing 4 x 15,000 and 4 x 75,000 cells respectively. We name the former “Very Low” and the latter “Low” chromatin pool. Next, we divide each chromatin pool into three equal aliquotes for immunoprecipitation against H3K4me3, H3K27ac and H3K27me3, while a small fraction of each pool (~ 1 µl) is set aside as Input control. This setup results in three ChIP reactions with 5000 cells/barcode for the “Very Low” pool and three ChIP reactions with 25,000 cells/barcode for the “Low” pool (Fig 3a).

**Figure 3:**
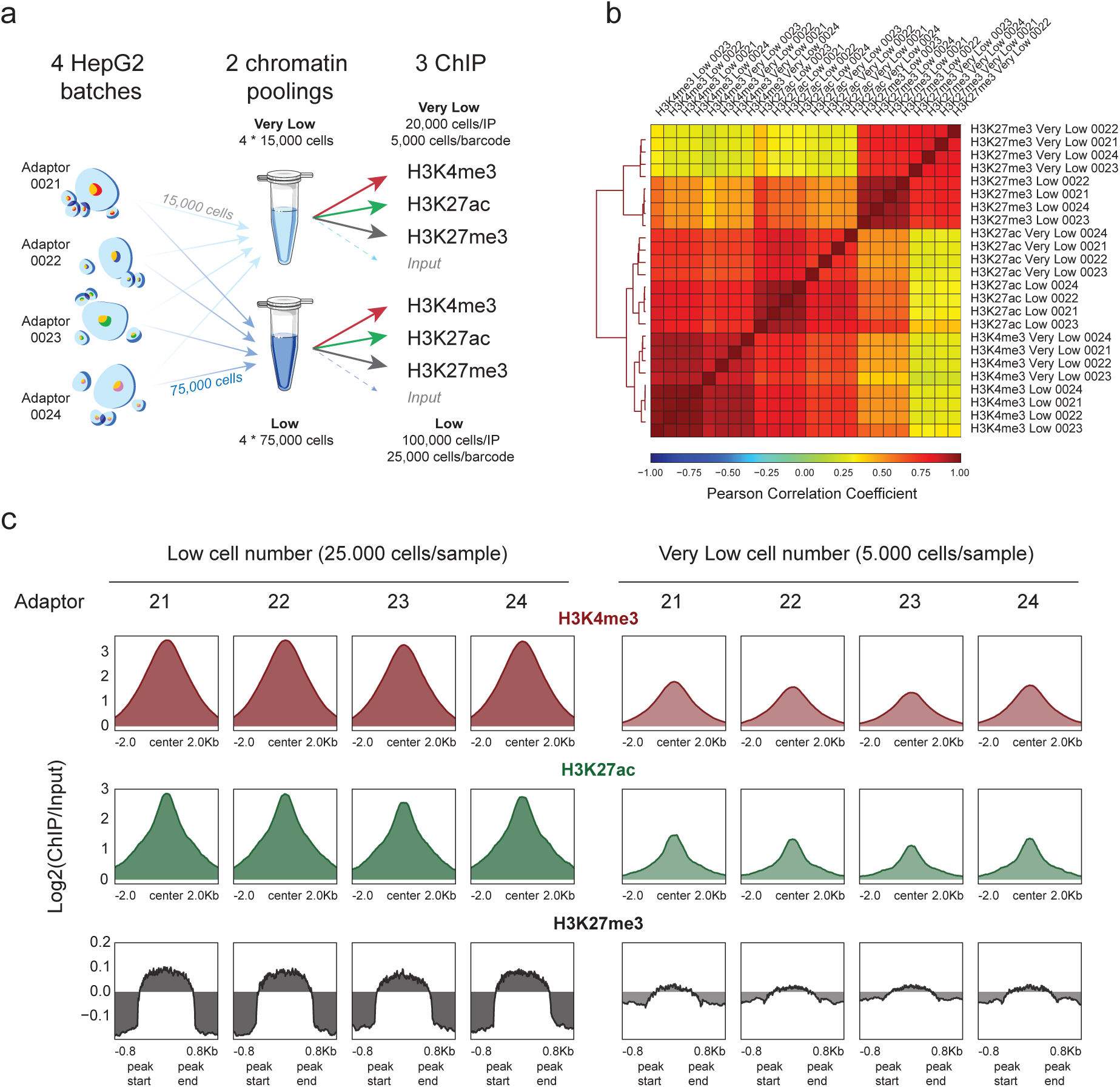
AutoRELACS works with low cell numbers. a) Overview of the experimental design used to test the sensitivity limits of AutoRELACS. Four batches of HepG2 cells are barcoded and pooled into two chromatin masterbatches, the first comprising 4 * 15,000 cells (Very Low input) and the second 4 * 75’000 cells (Low Input). Each chromatin pool is evenly split into three ChIP reactions (H3K4me3, H3K27ac, H3K27me3), while a small fraction (~ 1µl) is set aside as Input control. For the Very Low pool, about 20,000 cells are used in each ChIP, which corresponds to 5,000 cells/barcode. For the Low pool, about 100,000 cells are used in each ChIP, which corresponds to 25,000 cells/barcode. b) Hierarchical clustering of HepG2 ChIP-Seq profiles of H3K4me3, H3K27ac and H3K27me3, generated using Low and Very Low chromatin input, based on the pairwise Pearson Correlation Coefficient (PCC). Each pairwise PCC is computed based on the binned coverage (bin width = 10 kb) over the whole genome. c) Metaprofile of the mean enrichment over Input of H3K4me3 (upper panel, red), H3K27ac (central panel, green) and H3K27me3 (lower panel, grey), computed on a consensus set of peaks identified for each mark separately, from the Low and Very Low input chromatin.

The normalized genome-wide coverages coming from Low and Very Low experiments are highly correlated within histone modifications groups, which indicates that the profiles generated with different amounts of input material are overall similar (Fig 3b). Although we observe a deterioration of the signal-to-noise ratio in the Very Low group, the enrichment is preserved and, for narrow euchromatic marks, this is sufficient for robust peak calling (Fig 3c).

In summary, we show that AutoRELACS can be deployed for automated and parallelized profiling of protein-DNA interactions genome-wide also for limited amounts of biological samples.

## DISCUSSION

In this work we present AutoRELACS, an automated implementation of the RELACS protocol (9) that enables the automated high-throughput generation of ChIP-seq experiments. AutoRELACS natively interfaces with the computational pipelines offered by snakePipes (13), thus streamlining the generation and analysis of DNA-binding profiles at unprecedented scale.

RELACS can parallelize ChIP-seq data generation through *in situ* ligation of sample-specific barcodes into the digested chromatin inside the nuclear envelope. Here, we show that AutoRELACS successfully integrates a high number of barcodes in parallel, ensuring a balanced representation of each adaptor in the final chromatin pool. While we limit our test to 60 barcodes, a single AutoRELACS experiment can support the integration of up to 96 barcodes. The resulting chromatin pool can be split into 96 ChIP reactions, leading to the generation of up to 9,216 independent chromatin profiles in only three days. It should be noted that more imbalanced barcode distributions within the final chromatin pool may still lead to a successful profiling, at the cost of increasing the total sequencing depth. It is therefore suggested to perform a preliminary shallow sequencing of the chromatin input to estimate the total sequencing depth needed to ensure a minimum coverage for all samples.

We further show that AutoRELACS can generate high quality ChIP-seq data, comparable with the standards of the manual implementation, and that the method can be used for epigenetic profiling of low cell numbers. Together, these features suggest AutoRELACS as a method of choice in various clinical applications, potentially enabling comprehensive screening of epigenetic markers from small amounts of biological material.

The current AutoRELACS implementation has room for further improvements. To date, the method still requires human intervention in the earliest stages of the protocol. Future developments might integrate the use of focused sonicator platforms into the workflow of the liquid handler workstation, to further reduce user intervention and enable a full walk-away automated solution.

## MATERIALS AND METHODS

### Cell culture

S2 cells were cultured in Express Five SFM (Thermo Fisher Scientific) supplemented with glutamax, at 27°C and were provided by Akhtar’s lab (MPI-IE). HepG2 liver hepatocellular carcinoma (ATCC, HB-8065TM) were cultured in Eagle’s minimal essential medium (EMEM, Lonza, 06-174) supplemented with 10% fetal bovine serum (Sigma), 2 mM L-glutamine (Lonza), 1.8mM CaCl2, 1mM sodium pyruvate (Lonza) and penicillin–streptomycin mixture (100 units/mL, Lonza), at 37°C at 5% CO2 in 10cm plates, up to 70%-80% confluency.

### Cell fixation

HepG2 and S2 cells were fixed in 1% methanol-free formaldehyde (Thermo Scientific, 28906) in D-MEM (for HepG2 cells) or Express Five SFM (for S2 cells) for 15min at room temperature under gentle shaking. Formaldehyde was quenched for 5min by adding 125mM glycine final concentration. Cells were rinsed twice with ice-cold PBS, harvested by scraping (HepG2) and pelleted (300g, 10min, 4°C).

### Detailed AutoRELACS workflow

The AutoRELACS protocol is divided into five main steps (as described in Fig. 1).

A separated program file is provided for each automated section and is available for download at https://github.com/FrancescoFerrari88/AutoRELACS/tree/master/AutoRELACS_binaries_Biomek_i7.

#### 1) Nuclei extraction and chromatin digestion (manual protocol)

nuclei are extracted from fixed cells, swollen, digested, washed and counted as previously described (9). The resulting digested nuclei are resuspended in 10 mM Tris-HCl pH 8 at the nuclei density of 500,000 nuclei/25 µl (Drosophila S2) and 500,000 nuclei/25 µl (HepG2) for the following nuclei barcoding step.

#### 2) Chromatin barcoding and pooling (automated, method file “RELACS barcoding.bmf”)

in this step chromatin is barcoded inside the nuclei as previously described (9), but using automation. This method allows the processing for a flexible number of nuclei samples, from 1 to 96. Preparation of reagents: nuclei samples are aliquoted column-wise in a 96-wells PCR plate (25 µl of digested nuclei per well), named “Nuclei Plate”. 2 µl of the desired RELACS barcode at 15 µM are aliquoted in each well of a second 96-wells PCR plate, following the same coordinates of the respective nuclei aliquot (named “Index Plate”). The following reagent mixes are positioned into 1.5 ml conical tubes on the Biomek deck in a cold Peltier block: End Repair mix (ER), Ligation mix (LIG) and 3M NaCl, following directions as highlighted in the “guided instrument setup” (a screenshot of the deck is shown in Supplementary Fig. 2a).

Steps of the “RELACS barcoding” program: 5 µl of ER mix are added into each occupied well of “Nuclei Plate”. The plate is mixed on the orbital shaker present on the deck and incubated into the integrated PCR cycler for 30 min at 20 °C and for 5 min at 65 °C. End-repaired nuclei are transferred from “Nuclei Plate” to the “Index Plate” containing RELACS barcodes. 15.5 µl of LIG mix are added into each occupied well. The “IndexPlate” is shaken and transferred into the integrated PCR cycler for ligation incubation (15 min at 30 °C and for 15 min at 20 °C). The ligation is inactivated adding 5 µl of 3M NaCl into each occupied well of “Index Plate”. The plate is shaken and pooling is automatically performed by transferring samples from each occupied well of “Index Plate” to 1.5 ml tubes positioned into the “Final Pool” rack. Wells containing barcoded nuclei can be pooled as specified by the user, by indicating source and destination coordinates of “Index Plate” and “Final Pool” into the. csv file “Nuclei_Pooling_Template.csv”.

#### 3) Sonication-assisted nuclei lysis (manual protocol)

tubes containing nuclei pools are manually collected. Barcoded nuclei are pelletted down (5000 *g* for 10 min). Supernatants are discarded and pellets are resuspended into the desired volume of Shearing buffer supplemented with Protease Inhibitor Cocktail (Roche, 11873580001) and sonicated for 5 minutes in a Covaris E220 sonicator as described (9).

#### 4) ChIP and elution (automated, method file “RELACS ChIP-Elution.bmf”)

The method allows for a flexible number of ChIP reactions from 1 to 96 simultaneously. A screenshot of the overall organization of the deck is shown in Supplementary Fig. 2b.

All reagents used and the procedure of ChIP largely overlap to the ones described in our former publication (9), with the relevant modifications highlighted here below. Preparation of ChIP plate (named “Sample Plate”): ChIP reactions are carried out in a maximum volume of 150 µl instead of 200 µl used for manual RELACS. 75 µl of chromatin prepared in step 3 are aliquoted column-wise into a 1.2 ml storage plate (Thermo Fisher, AB1127) accordingly to the required number of ChIP. To equilibrate salts and detergents, 73 µl of 1X buffer iC1 (from iDeal ChIP-seq kit for histones, Diagenode C01010173) supplemented with Protease Inhibitor Cocktail (Roche, 11873580001) and 2 µl of 5M NaCl are added into each chromatin well. One µg per 100,000 cells of the desired antibody (H3K4me3 C15410003, H3K27ac C15410196, H3K27me3 C15410195, all from Diagenode) is added into each well. Remaining chromatins are set aside at 4 °C to prepare inputs. Please notice that input samples will be manually added later on before the automated decrosslinking step.

Preparation of reagents: ChIP Wash buffers 1 to 4 (from iDeal ChIP-seq kit for histones, Diagenode C01010173) are aliquoted into quarter module reservoirs divided by length. ChIP elution buffer (1% SDS, 200 mM NaCl, 10 mM Tris-HCl pH 8, 1 mM EDTA) is also aliquoted into the remaining well of the reservoir as highlighted in the “guided instrument setup”. ChIP beads (Dynabeads protein A-conjugated magnetic beads, Invitrogen) are washed twice with 1X buffer iC1 and aliquoted into two 1.5 ml conical tubes before placing them on the deck.

Automated protocol: the program involves four main steps (antibody incubation, beads incubation, ChIP washes, elution). Antibody incubation is performed by shaking the “Sample Plate” containing the ChIP reactions on the orbital shaker, repeating this procedure 12 times: 20 min continuous shaking at 500 rpm, stop for 10 min. In comparison to manual RELACS we carried out ChIP incubation for a total time of 6 hours at room temperature instead of 10 hours at 4 °C as used in manual RELACS. Please notice that we did this modification to overcome technical constraints that would have resulted in loss of samples when mixing by pipetting.

Beads incubation: beads placed on the deck are automatically mixed and 15 µl of beads are dispensed into each ChIP reaction. “Sample Plate” is then transferred on the orbital shaker and mixed for a total time of 2 hours at room temperature (5 min continuous shaking at 500 rpm, stop for 5 min, repeated 12 times). In comparison to the procedure used for manual RELACS, beads incubation time for AutoRELACS has been reduced by one hour.

ChIP washes: the following procedure is repeated for each of the four wash buffers. “Sample Plate” is transferred onto the magnetic rack and left for 5 minutes to reclaim the beads-bound immunocomplexes to the magnet. Supernatants are aspirated, discarded into the wash station, and 150 µl of wash buffer are added into each occupied well. Plate is shaken on the orbital shaker for about 5 minutes to wash the beads (5 seconds pulse shaking at 800 rpm for 60 times).

Elution: the last wash supernatants are removed from the beads. 80 µl of ChIP elution buffer is added to the beads and the plate is shaken on the orbital shaker for a total time of about 35 minutes (5 seconds pulse shaking at 800 rpm for 60 times, 4 minutes pause, for four times). “Sample Plate” is placed onto the magnet for 5 minutes and supernatants containing immunoprecipitated material are collected into a fresh 96-well plate (called “ChIP Eluates”) and stored overnight into the integrated PCR cycler at 10 °C.

#### 5) Decrosslink, purification, USER treatment, PCR amplification (automated, method file “RELACS Decrosslink-FinalLibrary.bmf”)

the plate “ChIP Eluates” is collected from the Biomek and Input samples are manually added column-wise after the ChIP samples (0.1-10% of the original chromatin volume in 80 µl of ChIP Elution buffer). This plate is placed back onto the deck and renamed in the instrument setup as “Sample Plate 2”.

Reagent preparation: 4 µl of 10 µM Illumina dual index primer cocktails (from IDT) are placed in a 96-well PCR plate column-wise following the desired pattern corresponding to the ChIP samples (plate is named “Index Plate”). The following reagents are required for this section of program, as specified in the instrument setup (Supplementary Fig. 2c): 100% isopropanol, EB (10 mM Tris-HCl pH 8), freshly prepared 85% ethanol (all on the deck at room temperature), proteinase K 20 mg/ml (Thermo Fisher, EO0491), glycogen 20 mg/mg (Thermo Fisher, R0561), carboxylated magnetic beads (Invitrogen, 65011), PCR mix (NEBNext Ultra II Q5 Master mix, NEB M0544), USER enzyme (NEB M5505), all placed in 1.5 ml conical tubes in a cold Peltier block. Ampure XP (Beckman Coulter, A63881) are thoroughly mixed and aliquoted column-wise according to the pattern of “Sample Plate 2” in a 96-well storage plate (AB0765, Thermo Fisher), using 100 µl of beads per well.

Automated Decrosslink: 2 µl of proteinase K are transferred into each occupied well of “Sample Plate 2” containing ChIP eluates and input samples. The plate is mixed on the orbital shaker and incubated for 2 hours at 65 °C into the integrated PCR cycler.

Automated DNA purification: in comparison to manual RELACS, in which decrosslinked DNA is purified using columns (Qiagen minElute PCR purification kit), AutoRELACS uses a custom-made DNA purification by precipitation and sequestration using carboxylated magnetic beads. Decrosslinked samples are transferred from the PCR plate to a larger 96-well storage plate (“ChIP Purification”, 4titude, LB0125). The following reagents are added into each occupied well: 2 µl of glycogen, 10 µl of carboxylated beads (automatically pre-mixed by pipetting before dispensing), and 80 µl of isopropanol. The plate “ChIP Purification” is mixed by shaking and incubated at room temperature for 10 minutes. The beads are reclaimed onto the integrated magnet for 5 minutes and supernatants are discarded. DNA bound to beads is washed twice using 200 µl of 85% ethanol. Beads are dried and DNA is automatically eluted by addition of 28 µl of EB into each occupied well. Plate is placed onto the magnet to discard the beads and to collect purified eluates.

USER treatment: 27 µl of purified DNAs are collected into a fresh 96-well PCR plate. 3 µl of USER enzyme is added into each occupied well. Plate is shaken and incubated into the integrated PCR cycler for 15 minutes at 37 °C. Samples are transferred into a 96-well storage plate for purification using Ampure XP (0.9X ratio). After purification, samples are eluted in 22 µl of EB. Automated amplification of final libraries and purification: 21 µl of each purified DNA are transferred to the 96-well PCR plate containing Illumina indexes (“Index Plate”). 25 µl of PCR mix are added into each occupied well and the plate is shaken. The plate is then transferred into the integrated PCR cycler for PCR incubation (hot start 98 °C for 30 sec; PCR cycles: 98 °C for 10 sec, 65 °C for 75 sec; final extension 65 °C for 5 min). Notice that before launching the method the user has the possibility of choosing the number of PCR cycles to use (10, 12 or 14). In the experiments presented in this work libraries were amplified using 12 PCR cycles (14 PCR cycles for low input ChIP). Amplified samples are transferred into a 96-well storage plate for double purification using Ampure XP (first at 0.8X ratio second at 1X ratio). Ready libraries are eluted in 25 µl of EB and transferred in a clean 96-well PCR plate.

### Sequencing

Libraries were quality-controlled to check the concentration (Qubit DNA HS, Invitrogen, Q32851) and the fragment size distribution (Fragment Analyzer capillary electrophoresis, NGS 1-6000 bp hs DNA kit). Libraries were pooled and normalized to 1 to 2 nM with 10% PhiX spike-in according to the Illumina guidelines. Libraries were clustered on NovaSeq XP flowcells and sequenced paired-end with a read length of 50 bp on an Illumina NovaSeq 6000 instrument.

### Bioinformatics analysis

BCL files were converted to fastq format using bcl2fastq2 (v. 2.20.0) and demultiplexed on illumina barcodes. Fastq files were used as input to snakePipes’ DNA-mapping and ChIP-seq workflows (v. 1.2.3) (13), using default parameters as listed in https://github.com/FrancescoFerrari88/AutoRELACS/tree/master/snakePipes_defaults. Mapping was performed on the genome build dm6 and hg38 for *D. melanogaster* and *H. sapiens* respectively. Briefly, fastq files were demultiplexed on RELACS adaptor barcodes and reads were mapped to the reference genome using Bowtie2 (v. 2.3) (15). Uniquely mapping read pairs (mapq > 3) were retained and duplicates were filtered on UMI using UMITools (paired mode) (v. 1.0.0) (16). Peaks were called using MACS2 (v. 2.1.2) (17) with default parameters. Merged peak sets were obtained by concatenating, sorting and merging peaks identified in the different experimental conditions included in the analysis, using bedtools sort | merge (v. 2.28) (18).

Clustered heatmaps, ChIP-seq metaprofiles and the clustered correlations heatmap were generated using deeptools (v. 3.3.1) (19), using filtered bam files as input. Principal component analysis (Fig 2a) was performed using the Python library scikit-learn (v. 0.19.1) on rlog-transformed count matrix (20). Coverage was obtained using deeptools’ multiBamSummary (v. 3.3.1) (19) on the merged peak set. We use pyGenomeTracks (21) to visualize signal tracks on specific genomic loci.

### Data and code availability

The fully reproducible and documented analysis is available on github at https://github.com/FrancescoFerrari88/AutoRELACS, as Jupyter notebooks and R/python scripts. Raw data and normalized bigWig tracks were deposited to GEO and are available for download using the following accession number: GSE147042.

## ACKNOWLEDGEMENTS

We would like to thank Alexiadis Anastasios for providing the S2 cells, and Erez Dror for technical support. This work was funded by the Deutsche Forschungsgemeinschaft - Project ID 192904750 - CRC 992 Medical Epigenetics.

## SUPPLEMENTARY

### Biomek i7 requirements and consumables for automation

The following instrument parts are required to perform AutoRELACS on the Biomek i7: Biomek i7 Workstation equipped with left and right pods; 1200 µl 96-multichannel (left pod), Span-8 pipets coupled with 1 ml syringe volume (right pod), gripper tools (one per pod), static Peltier with tube block for conical tubes, shaking Peltier and block for 96-well PCR plates, Orbital shaker, Wash station for multichannel, Wash station for Span-8, Magnet, Peristaltic pump (Masterflex L/S, Cole-Parmer), Automated PCR cycler (Thermo Fisher), seven Tip Loading Stations, twenty-six Automated Labware Positioners. Deck configuration details are indicated into each respective protocol part (Supplementary Fig. 2). Instrument configuration file is provided in the supplementary material (Biomeki7.bif).

The following plastic consumables are used for automation: Hard-Shell 96-well PCR plates (HSP9601, Bio-Rad), 96-Deep well storage microplates (4titude, LB0125), Low profile 1.2 ml square storage plate (AB1127, Thermo Fisher), 0.8 ml 96-well storage plate (AB0765, Thermo Fisher), Auto-sealing plate lids (MSL2022, Bio-Rad), Universal microplate lid (4ti-0290, 4titude), 300 ml reservoir (EK-2035, Agilent technologies), Modular reservoir quarter module divided by length (372788, Beckman Coulter), Modular reservoir quarter module (372790, Beckman Coulter), sterile tips with filter (all from Beckman Coulter): 1025 µl (B85955), 190 µl (B85911), 50 µl (B85888).

---

**Supplementary Figure 1:**
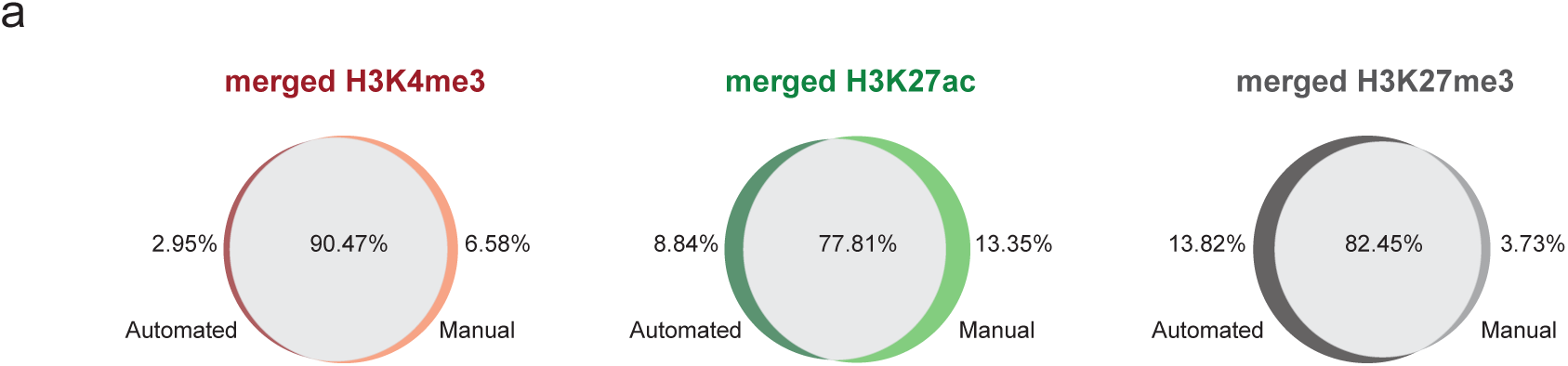
peaks identified in AutoRELACS and RELACS datasets overlap to a great extent. a) Venn diagrams representing the percentage of overlapping peaks and implementation-specific peaks identified in AutoRELACS (Automated) and manual RELACS (Manual) datasets, for H3K4me3 (left panel), H3K27ac (central panel) and H3K27me3 (right panel) profiles of S2 cells.

**Supplementary Figure 2:**
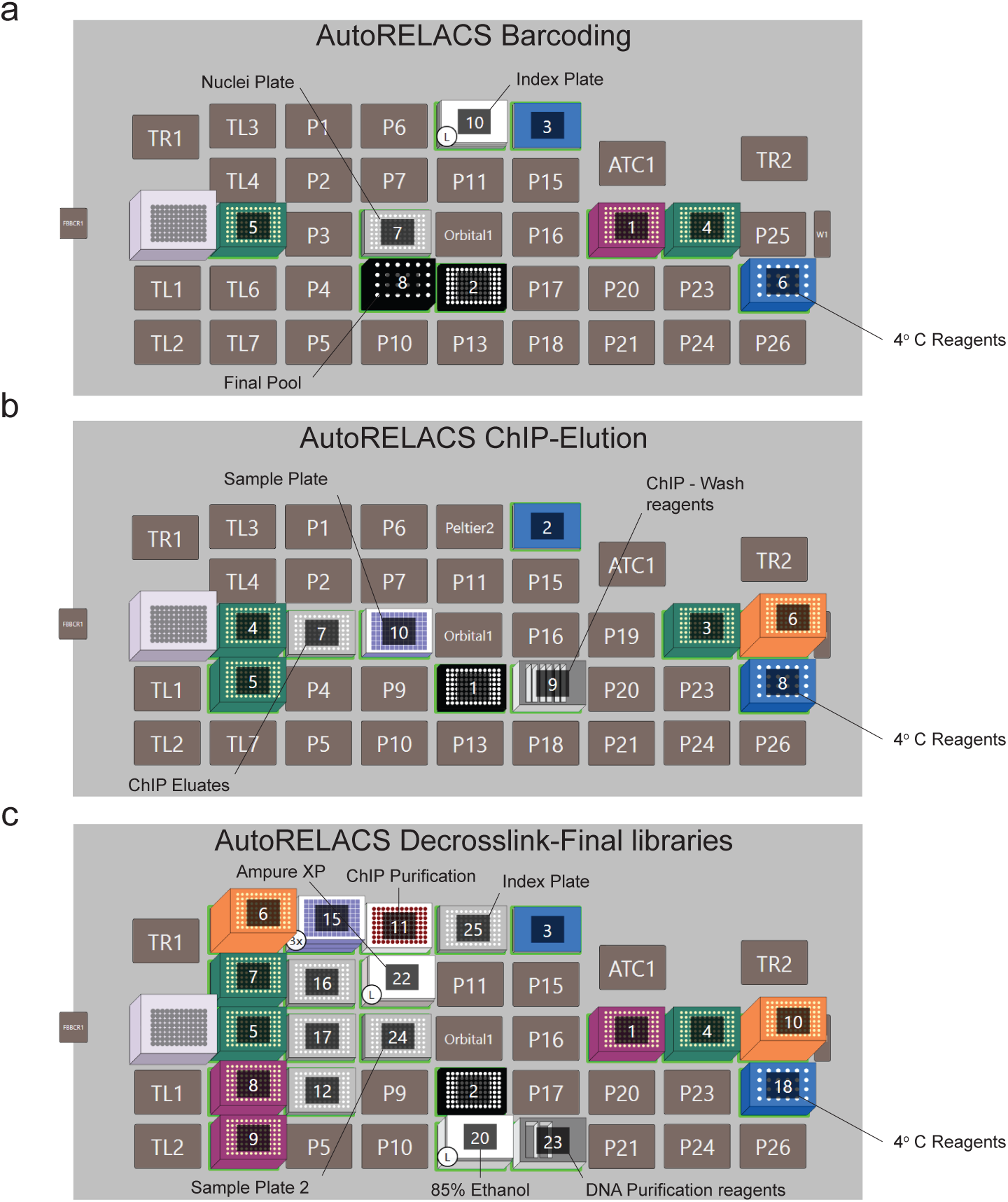
Biomek i7 deck configurations for AutoRELACS. a) Deck configuration for the method “RELACS barcoding”. On the deck are present filtered tips in different volumes (50 µl violet box in position 1, 190 µl green boxes in position 4 and 5), PCR lid for automation (3), magnet (2) and Peltier block containing barcoding reagents at 4 °C (4 °C reagents, 6). Digested nuclei are aliquoted in a 96-well PCR plate (Nuclei plate, 7). RELACS barcodes are aliquoted in a 96-well PCR plate (Index plate, 10) positioned on top of a cold Peltier. To protect the indexes, a plastic lid is positioned on top of the plate. b) Deck configuration for the method “RELACS ChIP-Elution”. On the deck are present filtered tips in different volumes (190 µl green boxes in position 3, 4 and 5, 1025 µl orange box in position 6), PCR lid for automation (2), magnet (1) and Peltier block containing ChIP reagents at 4 °C (4 °C reagents, 8). Room temperature ChIP reagents are stored in reservoirs (ChIP-Wash reagents, 9). ChIP reactions are aliquoted in a 96-deep well storage plate (Sample plate, 10). Final ChIP eluates are transferred into a 96-well PCR plate (ChIP eluates, 7). c) Deck configuration for the method “RELACS Decrosslink-FinalLibrary”. On the deck are present filtered tips in different volumes (50 µl violet boxes in position 1 8, 9, 190 µl green boxes in position 4, 5, 7, 1025 µl orange boxes in position 6, 10), PCR lid for automation (3), magnet (2) and Peltier block containing the required reagents at 4 °C (4 °C reagents, 18). Room temperature reagents are stored in reservoirs (DNA purification reagents, 23). 85% Ethanol is stored in a lidded reservoir (20). Ampure XP are aliquoted in a 96-deep well storage plate covered with a lid (Ampure XP, 22). ChIP and PCR purification occur in 96-deep well plates (11, 15). 96-well PCR plates in position 12, 16 and 17 are required for several steps of the method and to store the final libraries. ChIP and Input samples, which need to be firstly decrosslinked, are positioned in a 96-well PCR plate (24).

## Notes

https://github.com/FrancescoFerrari88/AutoRELACS

